# Cell-line specific role of Cathepsin B in triple-negative breast cancer growth, invasion and response to chemotherapy

**DOI:** 10.1101/2025.11.04.686610

**Authors:** Charlotte Kelley, Justinne R. Guarin, Emily Henrich, Jackson P. Fatherree, Claire M. Dunn, Anna Yui, Madeleine J. Oudin

## Abstract

Cathepsins are papain-family cysteine proteases known to play a cell-intrinsic role in protein degradation in the lysosome, as well as in digesting ECM and surface proteins after being secreted. Both of these functions are known to mediate pro-tumorigenic effects of CTSB in a range of cancers. Here, we specifically investigate the role of CTSB in TNBC, an aggressive subtype of breast cancer, where we find that high expression of CTSB in TNBC is associated with better outcomes. We used CRISPR to knockout CTSB in two highly metastatic TNBC cell lines, MDA-MB-231 and MDA-MB-468, and find different effects. In MDA-MB-231 cells, knockout of CTSB has no effect on cell viability, increases tumor cell 3D invasion in an ECM-independent manner, and increases sensitivity to many standard of care chemotherapy drugs. However, in MDA-MB-468 cells, knockout of CTSB increases cell viability, decreases tumor cell 3D invasion, in an ECM-independent manner, and drives resistance to certain chemotherapy drugs without affecting response to others. We find that in these cells, CTSB is not secreted, and that differential downstream mTOR and Akt activation can explain the differences seen in these phenotypes. Overall, our studies demonstrate that CTSB can regulate TNBC cell phenotypes via its lysosomal cell-intrinsic role, but that effects are cell-line specific, suggesting potential heterogeneity in the role of CTSB in TNBC.

## Introduction

Cathepsin B is part of the papain-family cysteine proteases, which are primarily located in the lysosome, and are involved in protein degradation, protein and lipid metabolism, autophagy and stress signaling ^1^. Cathepsins can also be secreted through lysosomal exocytosis, and extracellular CTSB is capable of directly digesting several extracellular matrix (ECM) proteins including collagen I, fibronectin, collagen IV, and laminin ^2 3^, cell surface proteins ^4^, and other proteases ^5^. Cathepsin B was first found to associated with cancer and metastatic potential in melanoma ^6^, localized in the lysosome of B16-F1 cells, as well as with increased activity on the cell membrane ^7^. Since, this work, cathepsin B has been linked to cancer progression and therapeutic responses in a number of cancers such as breast, colorectal, lung, ovarian, pancreatic cancer and osteosarcoma, and these studies suggest that the role of Cathepsin B in cancer progression is very context-dependent (reviewed in ^1^).

In breast cancer, several studies have shown upregulated levels of Cathepsin B levels in breast cancer patients compared to benign samples ^8,9^, with some studies showing that an association with earlier relapse and death ^10-12^. While none of these studies delve into specific subtypes of breast cancer, one study did find significant association with metastasis to the bone and poor disease survival, but this cohort had a majority of patients being ER, PR or Her2 positive ^13^. Triple-negative breast cancer (TNBC), which represents approximately 25% of breast cancer cases, is extremely hard to treat, due to its heterogeneous nature and the lack of targetable driver mutations^14^. Chemotherapy with anthracyclines and taxane-based drugs remains the most common treatment for patients with TNBC ^15, 16^. Pathologic complete response rates are low and vary between 15 to 40%. Hence, patients with TNBC have high rates of recurrence (over 30%), short time to recurrence, and poor survival ^17, 18^. Several clinical trials are ongoing to identify new agents to treat TNBC, targeting angiogenesis, DNA repair via PARP inhibitors, and cell proliferation via PI3K/Akt inhibitors ^19^. Recently, the PD-L1 inhibitor, atezolizumab, in combination with Abraxane was approved for patients with locally advanced or metastatic PD-L1–positive TNBC, however survival was increased by just 2.5 months ^20^. Understanding mechanisms of TNBC progression and chemoresistance could lead to the development of novel treatment strategies for TNBC, which could help improve patient outcome.

The function of CTSB in breast cancer progression has been investigated using PyMT-MMTV mice which spontaneously develop mammary tumors that metastasize to the lungs. Knockout of CTSB in these mice delays tumor growth, but does not affect metastasis, likely due to compensation from other proteases such as CTSZ ^21, 22^. Knockout of CTSB in PyMT cells decreased lung colonization in an experimental metastasis model, where cells are injected in the circulation ^21^. Further experiments showed that wild-type PyMT cells injected in a ctsb -/- host has reduced number and volume of lung metastases, demonstrating a role of stromal-expression of CTSB metastasis ^21^. While these clearly demonstrate an important role for CTSB in breast cancer progression, they do not investigate whether CTSB regulates invasion and metastasis via the lysosome or extracellular function. The role of CTSB in chemoresistance is not well known. Here, we set out to dissect the role of CTSB in driving tumor cell viability, invasion and response to chemotherapy in human TNBC. We investigated whether the ECM impacts CTSB-driven effects, and found that CTSB is not secreted by human TNBC cells, leading us to identify cell-line specific effects of lysosomal CTSB on downstream mTOR signaling in the context of TNBC.

## Results

### CTSB expression is high in TNBC and associated with better outcomes

While expression and prognostic value of CTSB has been widely studied in breast cancer cohorts, there have not been any studies that look at CTSB expression and association with outcome within individual breast cancer subtypes. We compared CTSB mRNA expression among breast cancer subtypes and found significantly higher levels of CTSB in TNBC patients compared to HER2 positive, and hormone positive cancers (Fig 1A). We looked at the association of CTSB expression levels with patient survival over time. As stated above, many studies have linked high CTSB levels with poor outcome in breast cancer cohorts that do not take subtype into account, and some that may be dominated by patients that are HER2 or hormone receptor positive. Here, we found that in the entire cohort, high CTSB mRNA levels are associated with significantly worse outcome, which is in line with previous work (Fig 1B). However, when we look at TNBC patients only, or basal breast cancer patients, high levels of CTSB are associated with significantly better outcomes (Fig 1C) ^23^. These data suggest that there may be unique and context-dependent roles for CTSB in breast cancer, specifically in

**Figure 1:**
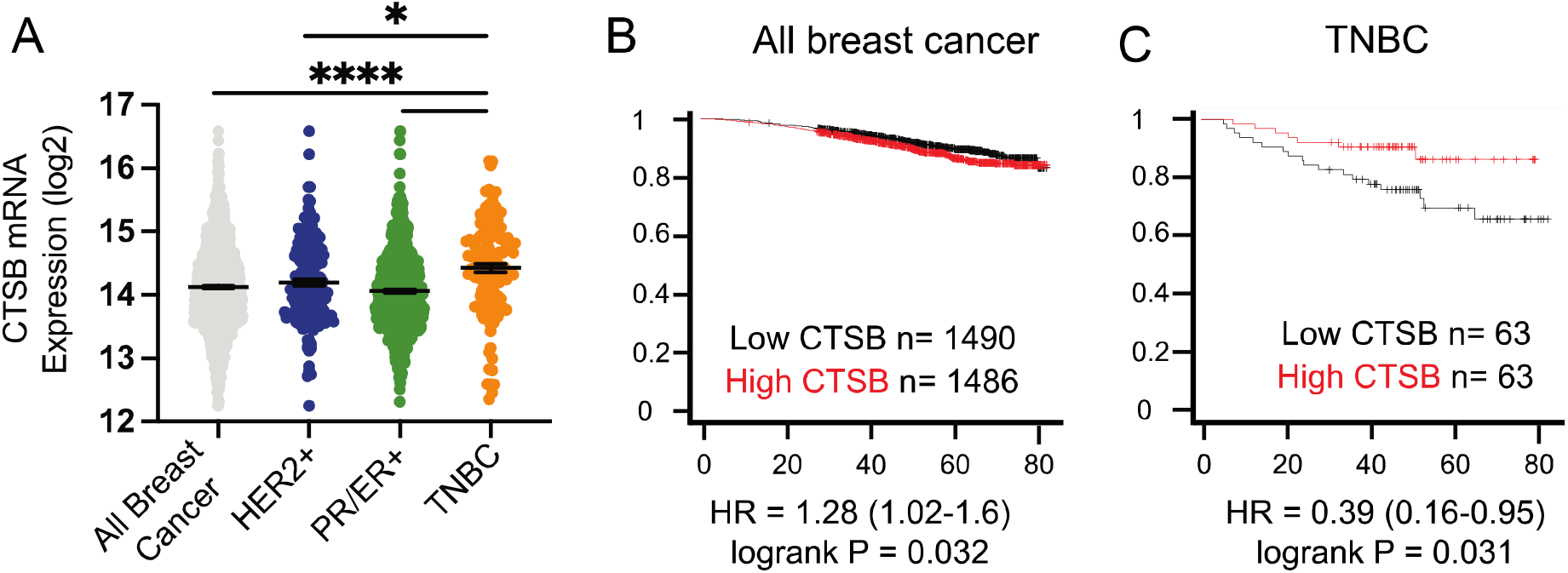
CTSB expression is correlated with improved patient outcomes in TNBC. (A) Patient data from cBioPortal for all breast cancer patients or aggregated by receptor subtype CTSB expression is increased in TNBC patients. High CTSB expression is correlated with poor patient outcome (B), however in TNBC high CTSB is associated with better patient outcome (C). data show mean ± SEM, * p<0.05, **p<0.01, ***p<0.005 by t-test.

### Knockdown of CTSB has cell-line specific effects on tumor cell growth and invasion

We then investigated the effect of CTSB levels on TNBC cell growth and invasion, phenotypes essential for tumor growth and invasion. We used CRISPR-Cas9 to generate knockdown of CTSB in two metastatic human TNBC cell lines: MDA-MB-231 cells, a cell line from a white American patient that is mesenchymal and MDA-MB-468, a cell line from an African American patient that is epithelial. We obtained over 90% knockdown in each cell line and both cell lines have similar level of CTSB protein expression (Fig 2A,D, Fig S1A,-C). To measure the role of CTSB on cell growth and viability, we seeded cells on plastic, Collagen I, Fibronectin or Collagen IV, ECM proteins that are known substrates of CTSB, can drive tumor cell invasion and are abundant in breast tumor tissue. Knockdown of CTSB in MDA-MB-231 cells did not affect cell proliferation, as measured by fold change in cell number over 72hr period, irrespective of the ECM substrate they were plated on (Fig 2B). We then used a 3D single cell invasion assay, where cells are seeded in 3D gels with Collagen I alone, or with Fibronectin or Collagen IV. Knockdown of CTSB in MDA-MB-231 cells increased tumor cell invasion speed and persistence in FN and Collagen IV (Fig 2B,C, S1D), suggesting that in these cells CTSB inhibits TNBC invasion. However, in MDA-MB-468 cells, we saw from different effects. Knockdown of CTSB in MDA-MB-468 significantly reduced cell proliferation, on plastic and on Collagen I, but the changes were not significant on Fibronectin or Collagen IV (Fig 2E). Further, knockdown of CTSB in MDA-MB-468 cells decreased tumor cell invasion speed Collagen I, FN and Collagen IV (Fig 2F), with no effects on persistence (Fig S1E). This shows that in MDA-MB-468 cells CTSB functions to promote cell growth and cellular invasion. Together, these data suggest that CTSB may be playing different roles in regulating metastasis across different TNBC cells.

**Figure 2:**
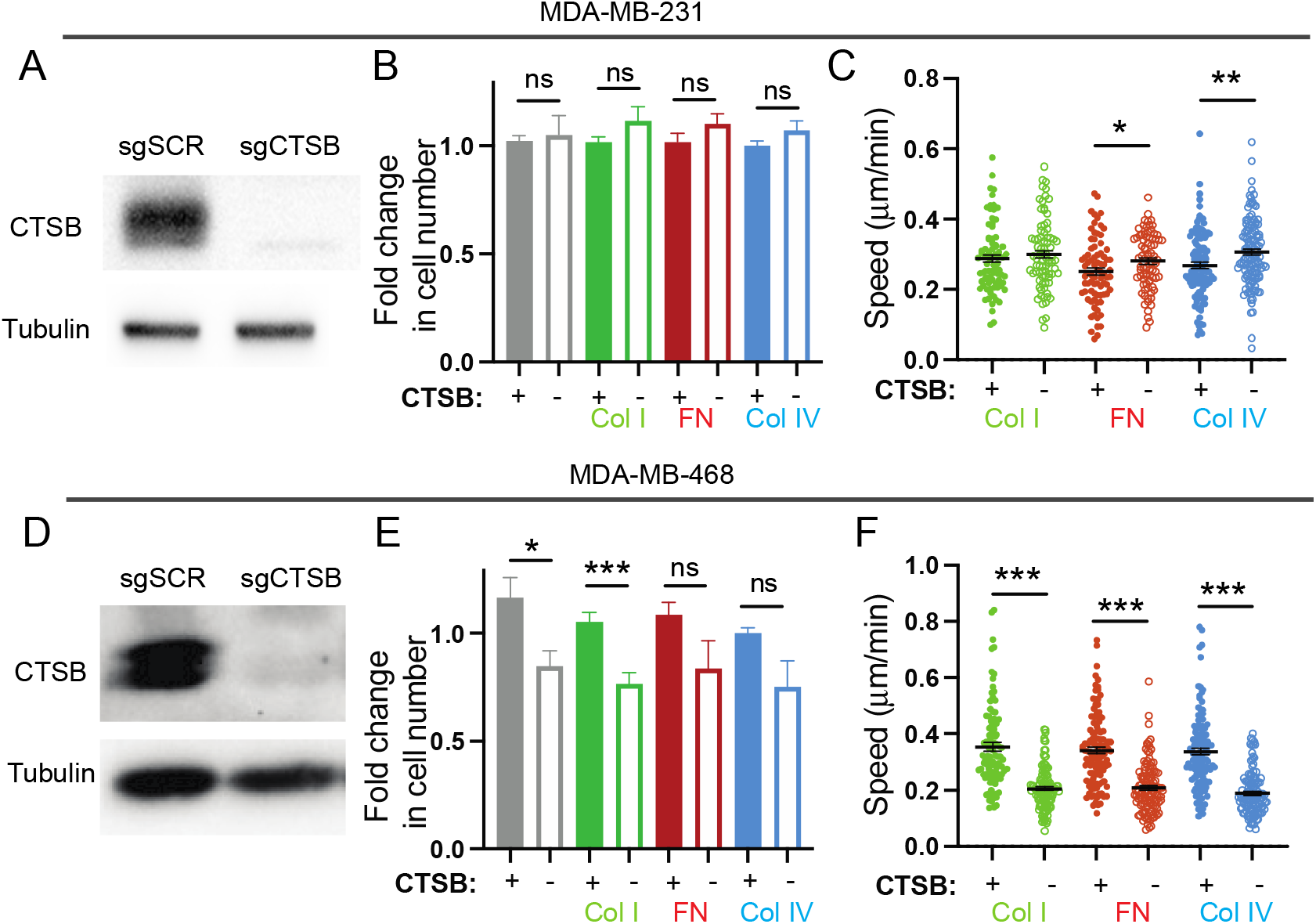
CTSB regulates tumor cell viability and invasion of TNBC cells. (A). Western Blot for CTSB protein in MDA-MB-231 cells with sgSCR and sgCTSB. B) Relative viability of MDA-MB-231 cells with sgSCR and sgCTSB plated on plastic, Collagen I (Col I), Fibronectin (FN) or Collagen IV(Col IV). C) 3D single cell invasion speed of MDA-MB-231 cells with sgSCR and sgCTSB seeded in hydrogels with Collagen I (Col I), Fibronectin (FN) or Collagen IV(Col IV). (D). Western Blot for CTSB protein in MDA-MB-468 cells with sgSCR and sgCTSB. E) Relative viability of MDA-MB-468 cells with sgSCR and sgCTSB plated on plastic, Collagen I (Col I), Fibronectin (FN) or Collagen IV(Col IV). F) 3D single cell invasion speed of MDA-MB-468 cells with sgSCR and sgCTSB seeded in hydrogels with Collagen I (Col I), Fibronectin (FN) or Collagen IV(Col IV). Data collected from at least 3 independent replicates, data show mean ± SEM, * p<0.05, **p<0.01, ***p<0.005 by t-test.

### Knockdown of CTSB has cell-line specific effects on chemotherapy response

We next examined the effect of CTSB on its role in promoting or sensitizing cells to chemotherapy. We used the four standard of care chemotherapy drugs used clinically to treat in TNBC: paclitaxel, doxorubicin, cisplatin and cyclophosphamide, and measured viability in cells with or without CTSB in the context of several ECM substrates when treated with chemotherapy was then measured. Relative viability was measured in CTSB knockout and control cells seeded on plastic, collagen I, collagen IV, or fibronectin and dosed with one of the four chemotherapies. Relative viability was normalized to the control (sgSCR) cells, thus if the relative viability is greater than one, the CTSB knockout cells are less sensitive to the drug. This increased resistance suggests that under normal conditions, CSTB functions to sensitize cells to chemotherapies. Conversely, if the knockout cells are more sensitive than the control cells, the relative viability is less than one, and thus CTSB functions to increase chemoresistance. In MDA-MB-231 cells, we found that knockdown of CTSB rendered cells more sensitive to multiple concentrations of paclitaxel and doxorubicin, independently of the ECM they were plated on (Fig 3A). The effect was less pronounced in response to Cisplatin, where effects were dependent on cells being plated on Collagen IV or FN. There are no consistent trends for cells treated with cyclophosphamide. However, in MDA-MB-468 cells, we found that knockdown of CTSB did not significantly impact response to paclitaxel, cisplatin or cyclophosphamide. Knockdown of CTSB did increase resistance to Doxorubicin, independently on the ECM protein it was plated on (Fig 3B). Overall, these data show that CTSB can play an important role in regulating response to chemotherapy, but these effects are cell line specific, and seem independent of the ECM environment.

**Figure 3:**
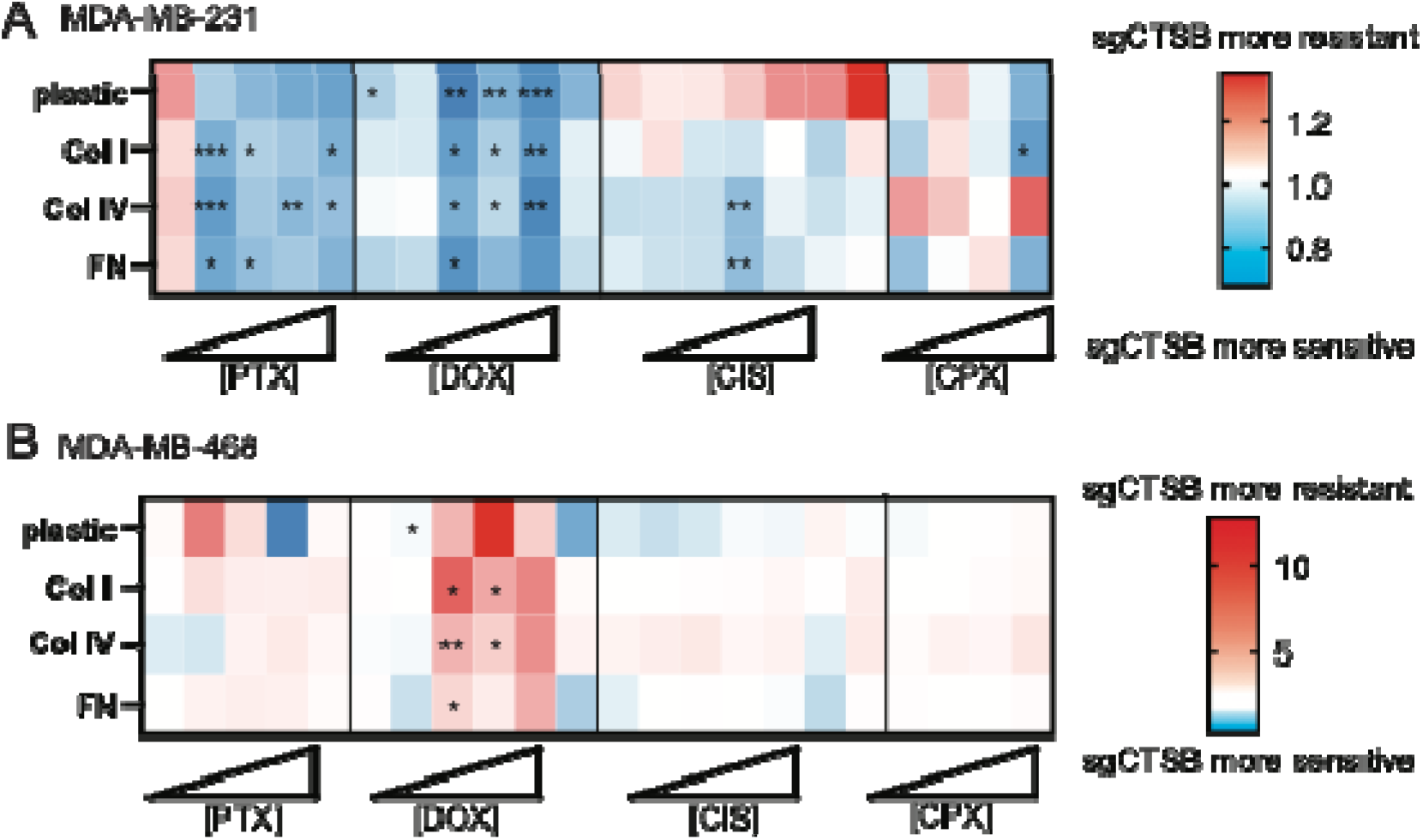
CTSB regula tes respo nse to chem other apy in some TNBC cells. Relati ve viabilit y of MDA-MB-231 (A) or MDA-MB-468 (B) cells with sgSCR or sgCTSB, with cells treated with Paclitaxel (1,10,100, 250, 1000nM), Doxorubicin (1,10,100, 250, 1000, 10000nM), Cisplatin (1,10,100nM, 2.5, 10 25uM), Cyclophosphamide (1, 2.5, 5, 10mM) and seeded on plastic, Collagen I, Collagen IV or Fibronectin. Data from at least 3 independent replicates, * p<0.05 **p<0.01, ***p<0.005 by t-test.

### CTSB is not secreted in human TNBC cells

We next investigated the mechanism by which CTSB may regulate tumor cell invasion and response to chemotherapy in TNBC cells. Cathepsin B is mainly located in the lysosome, but it can also be secreted through lysosomal exocytosis, where extracellular CTSB can digest ECM. We performed cell fractionation in MDA-MB-231 to investigate the localization of CTSB in the nucleus, cytoplasm, membrane and conditioned media. In both cell lines, we found that CTSB was not secreted into the media, but instead was confined to the membrane-bound and, to a lesser extent, the nuclear fractions (Fig 4A, B). We then investigated whether MDA-MB-468 cells secrete CTSB, and found that similarly, there was no CTSB found in the conditioned media obtained from MDA-MB-468 cells (Fig 4C). These data suggest that effects of CTSB in TNBC cells may not be due to direct effects on ECM digestion, but due rather to an intracellular function.

**Figure 4:**
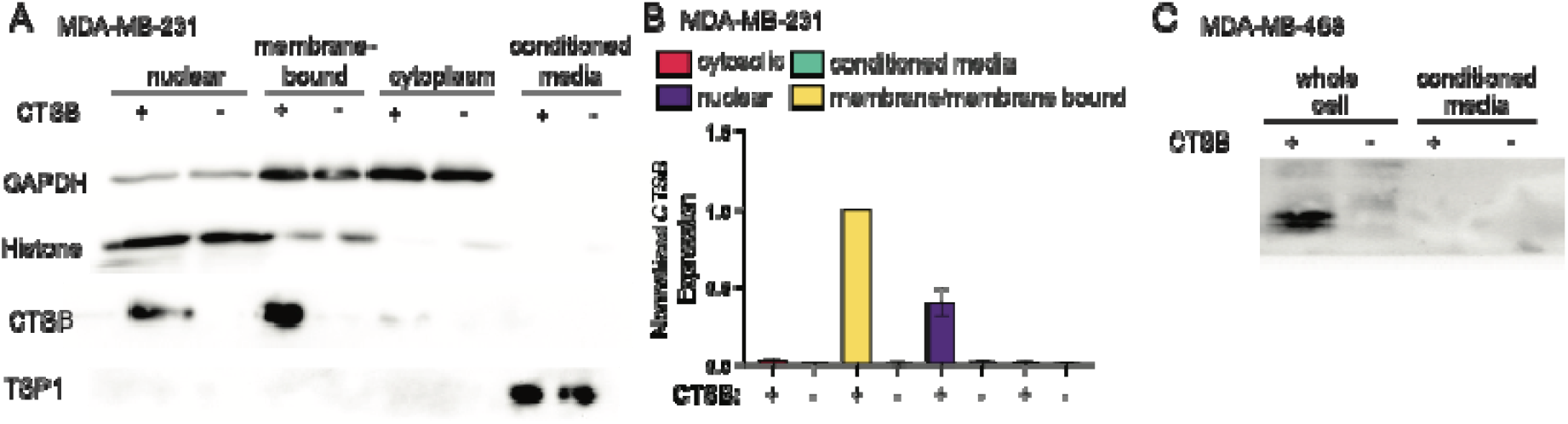
CTSB remains intracellular in TNBC cells. A) Western Blot for CTSB after cell fractionation in MDA-MB-231 cells to look at abundance in nuclear, membrane-bound, cytoplasmic and conditioned media. B) Quantification of CTSB abundance in different fractions. C) Western Blot for CTSB after cell fractionation in MDA-MB-468 cells to look at abundance in cells and conditioned media.

### mTOR signaling is regulated by CTSB

We next investigated differences in downstream signaling that may be driving these different effects on tumor cell growth, invasion and response to chemotherapy. We investigated activation of mTOR pathway by probing for phosphorylation at S2448, which is mostly mTORC1 and controls protein synthesis, cell growth and proliferation, and phosphorylation at S2481 which is mostly mTORC2 which regulates the actin cytoskeleton and promotes cell survival ^24^. In MDA-MB-231 cells, we found that knockdown of CTSB significantly increased phospho-mTOR at S2448, but had no effect at S2481, suggesting activation of mTROC1 (Fig 5A-C). Phospho-mTor S2448 is known to be phosphorylated by PI3K/AKT and we found increased AKT phosphorylation at S473 in MDA-MB-231 cells with CTSB knockdown (Fig 5D). However, we did not see an effect on mTOR signaling in MDA-MB-468 cells (Fig 5D-F). This suggests in MDA-MB-231 cells, there is enhanced AKT-mTOR signaling.

**Figure 5:**
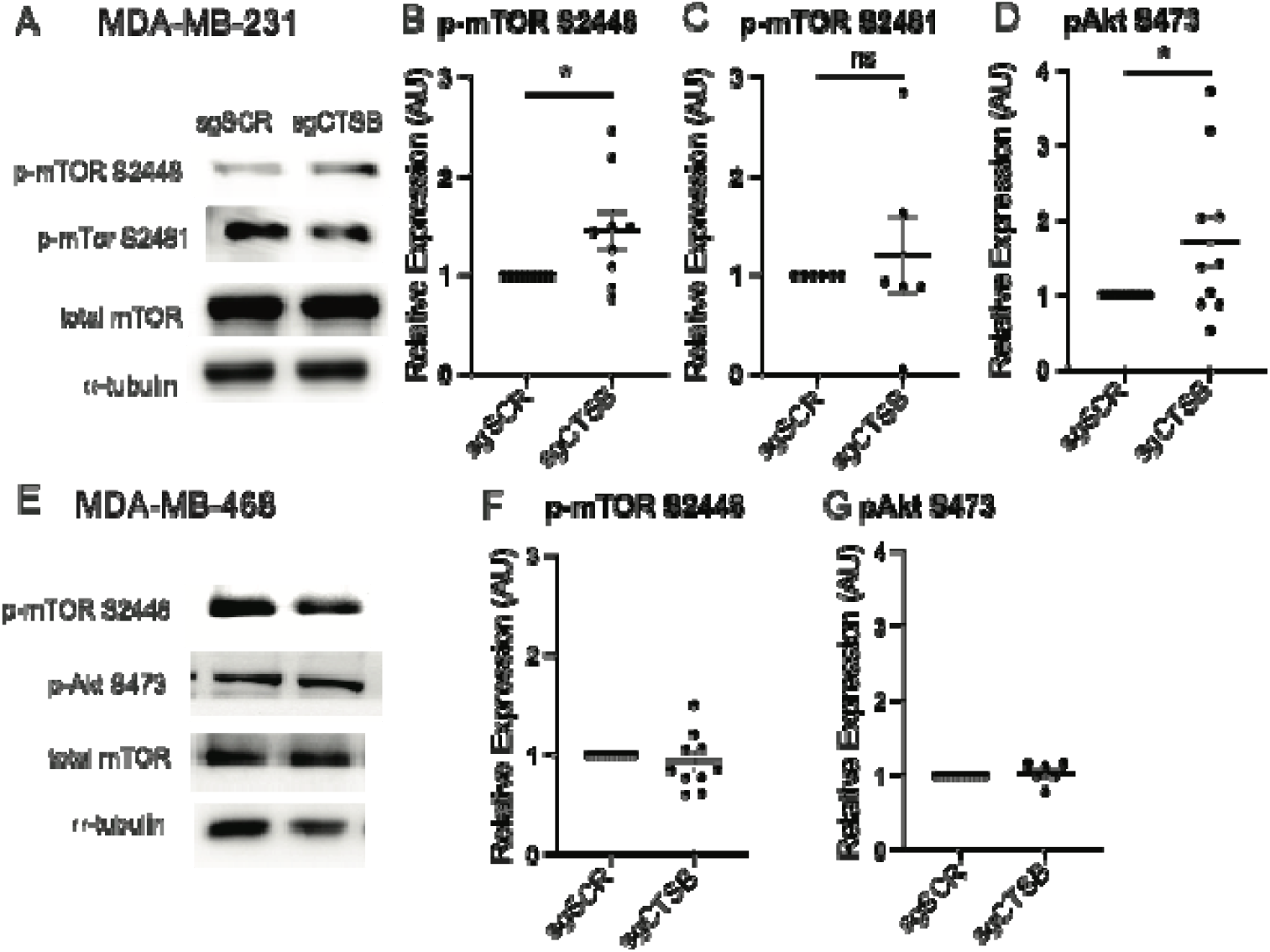
Knockdown of CTSB drives mTOR signaling in MDA-MB-231 cells but not in MDA-MB-468 cells. A) Representative Western Blot for MDA-MB-231 cells with sgSCR and sgCTSB for phosphorylated mTOR and total mTOR. Quantification of WB for p-mTOR S2448 (B)p-mTOR S2481 (C) and pAkt S473 (D) for MDA-MB-231 cells, normalized to total mTOR or AKT, respectively. E) Representative Western Blot for MDA-MB-468 cells with sgSCR and sgCTSB for phosphorylated mTOR and total mTOR. Quantification of WB for p-mTOR S2448 (F) and pAkt S473 (G) for MDA-MB-468 cells. Data from at least 5 biological replicates, data shown mean± SEM, * p<0.05 by t-test.

### Inhibition of mTOR signaling reverses effects of CTSB on tumor cell invasion and chemoresistance in MDA-MB-231 cells

We found that knockdown of CTSB increases phosphorylation of mTOR and Akt. Given that AKT-mTOR signaling regulates several downstream processes including protein translation, cell proliferation, autophagy, along with cell migration and associated cytoskeletal rearrangement, these changes likely leads to the increased invasion and increased sensitivity to certain chemotherapy drugs we see in the TNBC cells we tested. To further dissect the role of mTOR signaling in CTSB-driven effects on tumor cell invasion and chemoresistance, we used AZD3147, which is an extremely potent and selective dual inhibitor of mTORC1 and mTORC2. We first evaluated effects of mTOR inhibition on cell proliferation. After 24hrs of drug treatment, there were no effects on cell proliferation of control or sgCTSB MDA-MB-231 cells (Fig 6A). At 72hrs, we saw that mTOR inhibition with 10 nM AZD3147 significantly decreased tumor cell proliferation, independently of CTSB expression (Fig 6B). Next, we looked at invasion, where cells are treated for 24hrs and AZD3147 has no effects on cell survival. We found that inhibition of mTOR significantly decreased tumor cell invasion in CTSB knockout MDA-MB-231 cells, reversing the effect previously seen in the loss of CTSB (Fig 6C). Lastly, we looked at response to chemotherapy, which occurs over 72hr time frame. We found that inhibition of mTOR reversed the chemosensitivity induced by CTSB knockdown (Fig 6D). Together, these data show that inhibition of mTOR can reverse the effects of CTSB knockdown on tumor cell invasion and chemoresponse.

**Figure 6:**
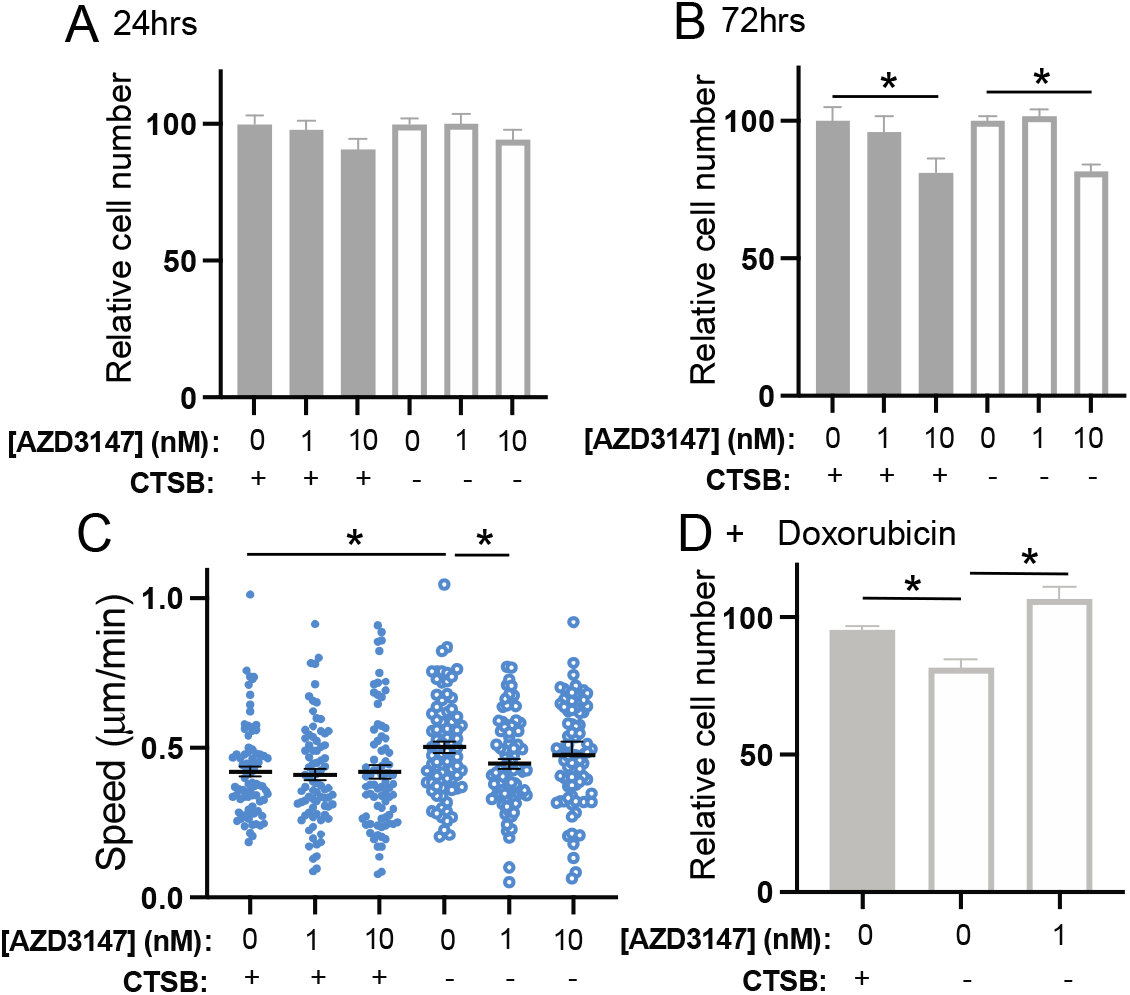
Inhibition of mTOR signaling reverses effects of CTSB on tumor cell invasion and chemoresistance in MDA-MB-231 cells. Quantification of cell viability by PrestoBlue assay after A) 24hrs and B)72hrs of AZD3147 treatment in MDA-MB-231 sgSCR and sgCTSB cells. C) Effect on invasion speed in 3D hydrogel with AZD3147 treatment. D) Quantification of cell viability with AZD3147 in combination with Doxorubicin. Data from at least 3 biological replicates, data show mean mean± SEM, * p<0.05, by t-test.

## Discussion

Cathepsins are papain-family cysteine proteases known to play a role cell intrinsically protein degradation in the lysosome, as well as in digesting ECM and surface proteins after being secreted. Both of these functions are known to mediate pro-tumorigenic effects of CTSB in a range of cancers. Here, we specifically investigate the role of CTSB in TNBC, an aggressive subtype of breast cancer, where we find that high expression of CTSB in TNBC is associated with better outcomes. We used CRISPR to knockout CTSB in two highly metastatic TNBC cell lines, MDA-MB-231 and MDA-MB-468, and find different effects. In MDA-MB-231 cells, knockout of CTSB has no effect on cell viability, increases tumor cell 3D invasion, in an ECM-independent manner, and increases sensitivity to many standard of care chemotherapy drugs. However, in MDA-MB-468 cells, knockout of CTSB increases cell viability, decreases tumor cell 3D invasion, in an ECM-independent manner, and drive resistance to many certain chemotherapy drugs, without affecting response to others. We find that in these cells, CTSB is not secreted, and that differential downstream mTOR and Akt activation can explain the differences seen in these phenotypes. Overall, our studies demonstrate that CTSB can regulate TNBC cell phenotypes via its lysosomal cell-intrinsic role, but that effects are cell-line specific, suggesting potential heterogeneity in the role of CTSB in TNBC.

Our data shows that CTSB is not secreted in TNBC cells *in vitro*, as cell fractionation shows that CTSB is not secreted from these cells and not detected in the conditioned media of both MDA-MB-231 and MDA-MB-468. CTSB is only found in the membrane-bound compartment, suggesting it is only in the lysosome. Previous studies in the field have not dissected this in detail, and all published work has focused on the PyMT-MMTV mouse model and PyMT-derived mouse cells. Specifically, *in vitro* studies in PyMT cells have shown that CTSB knockout alone does not affect 2D migration in scratch wound assay, but that is does reduce the number but not length of invasive strands in spheroid assays, and reduces invasion in a Matrigel transwell assay ^25^. While often associated with the lysosome, PyMT cells overexpressing CTSB increased matrix proteolysis as measured by dye-quenched Collagen IV, but this was not associated with a significant increase in tumor cell invasion in Collagen I ^2^, suggesting it may influence extracellular signaling. Consistent with this observation, specific inhibition of CTSB by CA074Me resulted in a decrease in invasion, indicating that enzymatic activity of CTSB, among other proteases, promotes tumor spheroid invasion into the matrix ^2^. Other studies have shown that transfection of MCF10A human breast epithelial cells with oncogenic *ras* leads to changes in the trafficking of cathepsin B, with increased membrane association ^26^, suggesting increased extracellular proteolytic activity. Further, CTSB was shown to colocalize with the annexin II tetramer on the cell surface of TNBC human breast cancer cells, which suggests it is released ^27^. However, none of these studies directly examined whether CTSB-driven effects are through direct effects on the ECM or via its intracellular function. Our data suggest that CTSB fuction may be influences by ECM and it may have indirect effects on the ECM via other proteases, but that its effects on cell viability, invasion and chemoresistance are driven by intracellular CTSB and downstream mTor and Akt activation.

Finally, our studies demonstrate cell-line specific effects of CTSB knockout. We used two human TNBC cell lines, which have previously shown behave similarly in a range of *in vitro* assays, and show high metastatic potential to lung and liver in immunocompromised mice ^28^. While both isolated from pleural effusions, they do differ in a number of characteristics: MDA-MB-231 cells are isolated from a Caucasian women, has low HER2 expression Kras and p53 mutation and is mesenchymal, MDA-MB-468 was isolated from an African American woman, it has no HER2 expression, PTEN deletion and EGFR amplification and is epithelial. African American women are 40 percent more likely to die from breast cancer than white women despite being diagnosed at similar rates. ^29^. There is evidence that average time to diagnosis is twice as long in African American women and surgery is less common, but also recent studies show that differential expression of pro-metastatic genes in tumors from African American patients relative to white patients ^30^. A significantly greater proportion of African American women were diagnosed with TNBC, and advanced (regional and distant) stages of breast cancer ^31^. Our studies demonstrate that it is important to perform mechanistic studies in a range of cell lines from patients from different backgrounds. It is imperative that we increase our understanding of mechanisms driving different outcomes among racial populations.

## Funding

This work was supported by NIH grant R01 to MJO.

## METHODS

### Antibodies and regents

Primary antibodies include: All antibodies were used at a concentration of 1:1000 unless otherwise noted. CTSB (ThermoFisher, 41-4800), alpha-tubulin (Cell Signaling Technology, 2144), GAPDH (14C10) (used at 1:2500, Cell Signaling Technology 50-190-704), histone (Cell Signaling Technology, 9715), Thrombospondin 1 (TSP) (ThermoFisher, MA5-13398), p-mTOR S2448 (Cell Signaling Technology, 2971), p-mTOR S2481 (Cell Signaling Technology, 2974), total mTOR (Cell Signaling Technology, 2983), p-AKT S473 (Cell Signaling Technology, 4060), or total AKT (Cell Signaling Technology, 9272).

Chemotherapeutics and inhibitors used were AZD3147 (ThermoFisher, 561510), Paclitaxel (Selleckchem, S1150), Doxorubicin (Selleckchem, E2516), and Cisplatin (Selleckchem, S1166). ECM proteins included Collagen I (Corning, 354236), native human Collagen IV (abcam, ab7536), Fibronectin (Sigma Aldrich, F1141)

### Cell Culture

MDA-MB-231 (ATCC, HTB-26) and MDA-MB-468 (ATCC, HTB-132) were cultured in Dulbecco’s Modified Eagle’s Medium (DMEM, Corning, MT10013CV) supplemented with 10% Fetal Bovine Serum (FBS, Gibco). Cells were regularly assessed for mycoplasma using the Universal Mycoplasma Detection Kit (ATCC, 30-1012 K). Knockouts were generated by first generating MDA-MB-231 or MDA-MB-468 stably expressing Cas9 cells through lentiviral transfection. Next, guides were transfected from the LentiArray(TM) Human Protease CRISPR Library (ThermoFisher, A42279). Knockout was confirmed using western blot. All cells were cultured at 37 C and 5% CO2.

### Human expression data

Human expression data is obtained from cBioportal, using publicly available datasets from ^32 33^.

### Cell viability

5,000 cells were seeded on either plastic tissue-culture grade 96-well plates or plates pre-coated with 20 ug/ml of either collagen I, collagen IV, or fibronectin. Cells were allowed to adhere for 24 hours, then viability was measured using the PrestoBlue Cell Viability Reagent (Invitrogen, A13261) to quantify the relative cell viability on day 1. Cells were then incubated for an additional 72 hours at 37C, 5% CO2 and viability was remeasured. Results are plotted as a change in viability relative to the control sgSCR cells.

### 3D Cell invasion

Assays performed as previously published^28^. 20,000 cells were suspended in media plus an ECM mixture of 1 mg/ml collagen I with or without one of the following: 50 ug/ml fibronectin or 20 ug/ml collagen IV in a 48-well plate and allowed to incubate at 37 C at 5% CO2. After two hours, 50 ul of cell culture media was added and if treated with inhibitors, these were added in this step. Wells were imaged using a Keyence BZ-X710 microscope (Keyence) for 16-hours at 10-minute intervals while being kept in a humidified, 37C, 5% CO2 environment. Cell movements were tracked using the VQ-9000 Video Editing/Analysis Software (Keyence) and invasive speed and persistence were calculated using our custom Matlab script (MathWorks).

### Western Blotting

Protein lysates separated on a 4-12% SDS-PAGE polyacrylamide gel and transferred to a nitrocellulose membrane using the TransBlot Turbo Transfer system (Biorad). Next, membranes were blocked in 5% nonfat dry milk resolubilized in trie-buffered saline and 0.05% Tween-20 (TBST) before being incubated in primary antibodies overnight in 1% nonfat dry milk in TBST. After three washes in TBS-T, secondary antibodies conjugated with HRP were incubated, washed again in TBST, then imaged using XX reagent and a ChemiDoc MP imager (Biorad, 12003154).

### Cell Fractionation

Cells were grown in a 10-cm dish in 10 ml of appropriate media for 48 hours. This conditioned media was first treated with a protease and phosphatase inhibitors and then concentrated using a 3 MWCO spin column (ThermoFisher, 88525). Remaining cells were washed in PBS, then collected using Trypsin. 5 ml media containing 10% FBS was used to inactivate remaining trypsin, then cells were pelleted and sequentially lysed then spun following published protocols^34^. Cellular fractions were run on SDS PAGE gels and purity was assessed using appropriate antibodies for given cellular compartments.

### Immunostaining

Cells were fixed using 4% PFA for 10 min. Cells were permeabilized with 0.2% triton-X washed and then blocked with 5% normal donkey serum, and 0.1% tween-20 overnight at 4°C. Primary antibodies were then added in a 1% normal donkey serum and 0.1% triton x solution and incubated overnight at 4°C. Secondaries including Phalloidin (F-actin) and DAPI (nuclei) were then added in a 1% normal donkey serum and 0.1% triton x solution. Cells were then washed and imaged using a Keyence BZ-X710 microscope (Keyence, Elmwood Park, NJ).

### Statistical analysis

GraphPad Prism v9.1.0 was used for the generation of graphs and statistical analysis. Significance was determined using a one-way ANOVA with Dunnett’s multiple comparisons test unless otherwise stated. MATLAB was used to determine linear relationships and correlation.

## Funding

This work was funded by R01CA255742 to MJO from NCI, as well as funds from the Tufts School of Engineering start-up funds to MJO.

## Supplemental Figures

**Figure S1:**
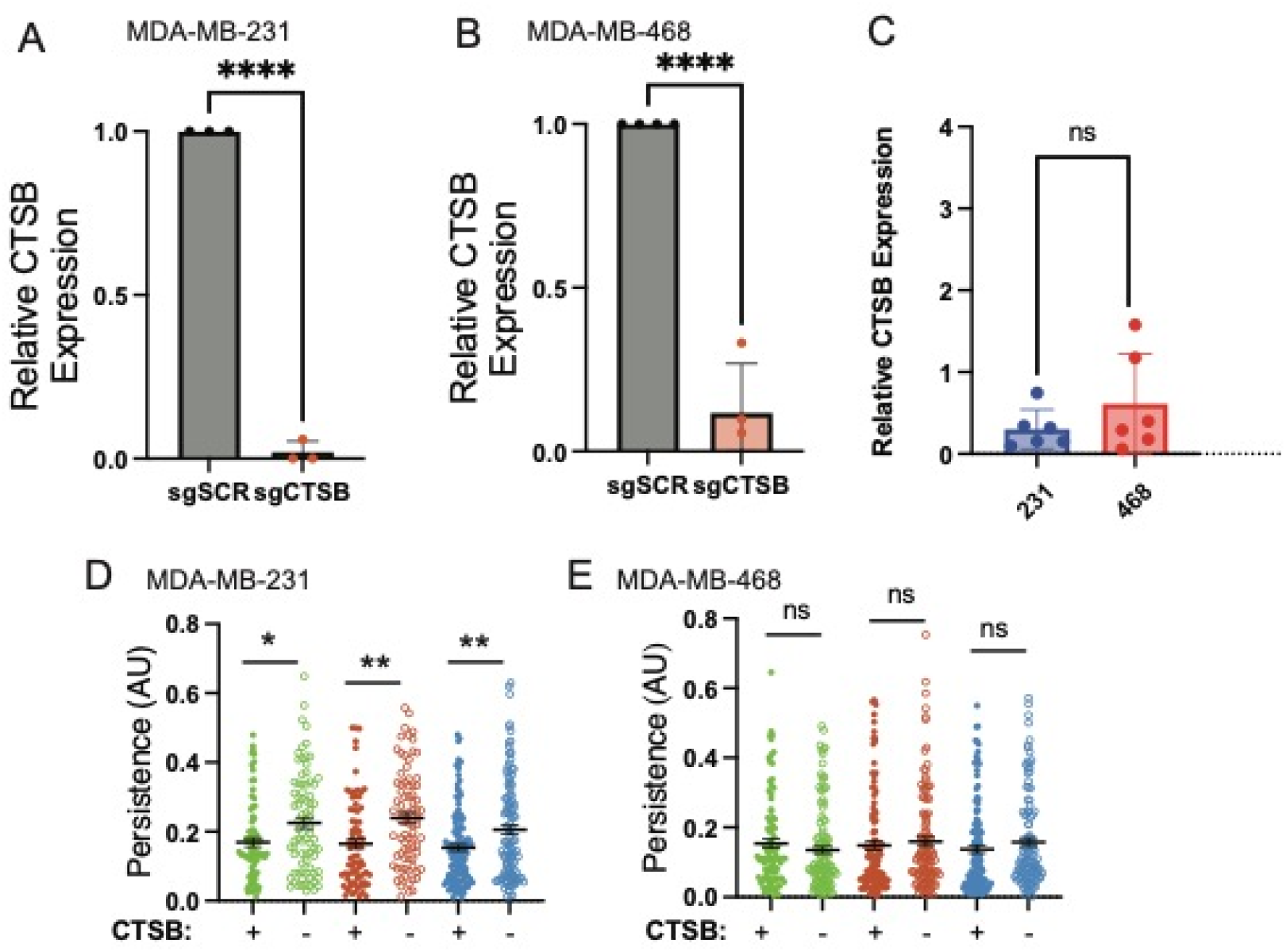
CTSB knockdown in TNBC cells. Quantification of Western Blots from CTSB levels in sgSCR and sgCTSB in MDA-MB-231 (A) and MDA-MB-468 cells (B). Comparison of normalized CTSB protein levels in MDA-MB-231 and MDA-MB-468 cells (C). 3D single cell persistence of D) MDA-MB-231 and E) MDA-MB-468 cells with sgSCR and sgCTSB seeded in hydrogels with Collagen I (Col I), Fibronectin (FN) or Collagen IV(Col IV). Data collected from at least 3 independent replicates, data show mean± SEM, * p<0.05, **p<0.01, ***p<0.005 by t-test.

## Notes

### Competing Interest Statement

The authors have declared no competing interest.

## References

1. Olson OC, Joyce JA. Cysteine cathepsin proteases: regulators of cancer progression and therapeutic response. Nat Rev Cancer. 2015;15(12):712–29. doi: 10.1038/nrc4027. PubMed PMID: 26597527.

2. Bengsch F, Buck A, Gunther SC, Seiz JR, Tacke M, Pfeifer D, von Elverfeldt D, Sevenich L, Hillebrand LE, Kern U, Sameni M, Peters C, Sloane BF, Reinheckel T. Cell type-dependent pathogenic functions of overexpressed human cathepsin B in murine breast cancer progression. Oncogene. 2014;33(36):4474–84. Epub 20130930. doi: 10.1038/onc.2013.395. PubMed PMID: 24077280; PMCID: PMC4139469.

3. Schmitz J, Gilberg E, Loser R, Bajorath J, Bartz U, Gutschow M. Cathepsin B: Active site mapping with peptidic substrates and inhibitors. Bioorg Med Chem. 2019;27(1):1–15. Epub 20181019. doi: 10.1016/j.bmc.2018.10.017. PubMed PMID: 30473362.

4. Mizunoe Y, Kobayashi M, Hoshino S, Tagawa R, Itagawa R, Hoshino A, Okita N, Sudo Y, Nakagawa Y, Shimano H, Higami Y. Cathepsin B overexpression induces degradation of perilipin 1 to cause lipid metabolism dysfunction in adipocytes. Sci Rep. 2020;10(1):634. Epub 20200120. doi: 10.1038/s41598-020-57428-6. PubMed PMID: 31959889; PMCID: PMC6971249.

5. Kobayashi H, Moniwa N, Sugimura M, Shinohara H, Ohi H, Terao T. Effects of membrane-associated cathepsin B on the activation of receptor-bound prourokinase and subsequent invasion of reconstituted basement membranes. Biochim Biophys Acta. 1993;1178(1):55–62. doi: 10.1016/0167-4889(93)90109-3. PubMed PMID: 8329457.

6. Sloane BF, Dunn JR, Honn KV. Lysosomal cathepsin B: correlation with metastatic potential. Science. 1981;212(4499):1151–3. doi: 10.1126/science.7233209. PubMed PMID: 7233209.

7. Sloane BF, Rozhin J, Johnson K, Taylor H, Crissman JD, Honn KV. Cathepsin B: association with plasma membrane in metastatic tumors. Proc Natl Acad Sci U S A. 1986;83(8):2483–7. doi: 10.1073/pnas.83.8.2483. PubMed PMID: 3458210; PMCID: PMC323322.

8. Thomssen C, Schmitt M, Goretzki L, Oppelt P, Pache L, Dettmar P, Janicke F, Graeff H. Prognostic value of the cysteine proteases cathepsins B and cathepsin L in human breast cancer. Clin Cancer Res. 1995;1(7):741-6. PubMed PMID: 9816040.

9. Nouh MA, Mohamed MM, El-Shinawi M, Shaalan MA, Cavallo-Medved D, Khaled HM, Sloane BF. Cathepsin B: a potential prognostic marker for inflammatory breast cancer. J Transl Med. 2011;9:1. Epub 20110103. doi: 10.1186/1479-5876-9-1. PubMed PMID: 21199580; PMCID: PMC3022726.

10. Foekens JA, Kos J, Peters HA, Krasovec M, Look MP, Cimerman N, Meijer-van Gelder ME, Henzen-Logmans SC, van Putten WL, Klijn JG. Prognostic significance of cathepsins B and L in primary human breast cancer. J Clin Oncol. 1998;16(3):1013–21. doi: 10.1200/JCO.1998.16.3.1013. PubMed PMID: 9508185.

11. Lah TT, Cercek M, Blejec A, Kos J, Gorodetsky E, Somers R, Daskal I. Cathepsin B, a prognostic indicator in lymph node-negative breast carcinoma patients: comparison with cathepsin D, cathepsin L, and other clinical indicators. Clin Cancer Res. 2000;6(2):578-84. PubMed PMID: 10690542.

12. Lah TT, Kalman E, Najjar D, Gorodetsky E, Brennan P, Somers R, Daskal I. Cells producing cathepsins D, B, and L in human breast carcinoma and their association with prognosis. Hum Pathol. 2000;31(2):149–60. doi: 10.1016/s0046-8177(00)80214-2. PubMed PMID: 10685628.

13. Sun T, Jiang D, Zhang L, Su Q, Mao W, Jiang C. Expression profile of cathepsins indicates the potential of cathepsins B and D as prognostic factors in breast cancer patients. Oncol Lett. 2016;11(1):575–83. Epub 2016/02/13. doi: 10.3892/ol.2015.3960. PubMed PMID: 26870250; PMCID: PMC4727043.

14. Triple-negative breast cancer: Challenges and opportunities of a heterogeneous disease, (2016).

15. Al-Mahmood S, Sapiezynski J, Garbuzenko OB, Minko T. Metastatic and triple-negative breast cancer: challenges and treatment options. Drug Deliv Transl Res. 2018;8(5):1483–507. Epub 2018/07/07. doi: 10.1007/s13346-018-0551-3. PubMed PMID: 29978332; PMCID: PMC6133085.

16. Sharma P. Update on the Treatment of Early-Stage Triple-Negative Breast Cancer. Curr Treat Options Oncol. 2018;19(5):22. Epub 2018/04/16. doi: 10.1007/s11864-018-0539-8. PubMed PMID: 29656345.

17. Li CH, Karantza V, Aktan G, Lala M. Current treatment landscape for patients with locally recurrent inoperable or metastatic triple-negative breast cancer: a systematic literature review. Breast cancer research : BCR. 2019;21(1):143. Epub 2019/12/18. doi: 10.1186/s13058-019-1210-4. PubMed PMID: 31842957; PMCID: PMC6916124.

18. Cortazar P, Zhang L, Untch M, Mehta K, Costantino JP, Wolmark N, Bonnefoi H, Cameron D, Gianni L, Valagussa P, Swain SM, Prowell T, Loibl S, Wickerham DL, Bogaerts J, Baselga J, Perou C, Blumenthal G, Blohmer J, Mamounas EP, Bergh J, Semiglazov V, Justice R, Eidtmann H, Paik S, Piccart M, Sridhara R, Fasching PA, Slaets L, Tang S, Gerber B, Geyer CE, Jr., Pazdur R, Ditsch N, Rastogi P, Eiermann W, von Minckwitz G. Pathological complete response and long-term clinical benefit in breast cancer: the CTNeoBC pooled analysis. Lancet. 2014;384(9938):164–72. Epub 2014/02/18. doi: 10.1016/S0140-6736(13)62422-8. PubMed PMID: 24529560.

19. Bergin ART, Loi S. Triple-negative breast cancer: recent treatment advances. F1000Res. 2019;8. Epub 2019/08/27. doi: 10.12688/f1000research.18888.1. PubMed PMID: 31448088; PMCID: PMC6681627.

20. Schmid P, Adams S, Rugo HS, Schneeweiss A, Barrios CH, Iwata H, Dieras V, Hegg R, Im SA, Shaw Wright G, Henschel V, Molinero L, Chui SY, Funke R, Husain A, Winer EP, Loi S, Emens LA, Investigators IMT. Atezolizumab and Nab-Paclitaxel in Advanced Triple-Negative Breast Cancer. N Engl J Med. 2018;379(22):2108–21. Epub 2018/10/23. doi: 10.1056/NEJMoa1809615. PubMed PMID: 30345906.

21. Vasiljeva O, Papazoglou A, Kruger A, Brodoefel H, Korovin M, Deussing J, Augustin N, Nielsen BS, Almholt K, Bogyo M, Peters C, Reinheckel T. Tumor cell-derived and macrophage-derived cathepsin B promotes progression and lung metastasis of mammary cancer. Cancer Res. 2006;66(10):5242–50. doi: 10.1158/0008-5472.CAN-05-4463. PubMed PMID: 16707449.

22. Sevenich L, Werner F, Gajda M, Schurigt U, Sieber C, Muller S, Follo M, Peters C, Reinheckel T. Transgenic expression of human cathepsin B promotes progression and metastasis of polyoma-middle-T-induced breast cancer in mice. Oncogene. 2011;30(1):54–64. Epub 20100906. doi: 10.1038/onc.2010.387. PubMed PMID: 20818432.

23. Gyorffy B. Survival analysis across the entire transcriptome identifies biomarkers with the highest prognostic power in breast cancer. Comput Struct Biotechnol J. 2021;19:4101-9. Epub 20210718. doi: 10.1016/j.csbj.2021.07.014. PubMed PMID: 34527184; PMCID: PMC8339292.

24. Watanabe R, Wei L, Huang J. mTOR signaling, function, novel inhibitors, and therapeutic targets. J Nucl Med. 2011;52(4):497–500. Epub 20110318. doi: 10.2967/jnumed.111.089623. PubMed PMID: 21421716.

25. Sevenich L, Schurigt U, Sachse K, Gajda M, Werner F, Muller S, Vasiljeva O, Schwinde A, Klemm N, Deussing J, Peters C, Reinheckel T. Synergistic antitumor effects of combined cathepsin B and cathepsin Z deficiencies on breast cancer progression and metastasis in mice. Proc Natl Acad Sci U S A. 2010;107(6):2497-502. Epub 20100121. doi: 10.1073/pnas.0907240107. PubMed PMID: 20133781; PMCID: PMC2823914.

26. Sloane BF, Moin K, Sameni M, Tait LR, Rozhin J, Ziegler G. Membrane association of cathepsin B can be induced by transfection of human breast epithelial cells with c-Ha-ras oncogene. J Cell Sci. 1994;107 (Pt 2):373–84. doi: 10.1242/jcs.107.2.373. PubMed PMID: 8207069.

27. Mai J, Finley RL, Jr., Waisman DM, Sloane BF. Human procathepsin B interacts with the annexin II tetramer on the surface of tumor cells. J Biol Chem. 2000;275(17):12806–12. doi: 10.1074/jbc.275.17.12806. PubMed PMID: 10777578.

28. Conner SJ, Guarin JR, L. TT, Fatherree JP, Kelley C, Payne SL, Parker SR, Bloomer H, Zhang C, Salhany K, McGinn RA, Henrich E, Yui A, Srinivasan D, Borges H, Oudin MJ. Cell morphology best predicts tumorigenicity and metastasis in vivo across multiple TNBC cell lines of different metastatic potential. Breast cancer research : BCR. 2024;26(1):43. Epub 20240311. doi: 10.1186/s13058-024-01796-8. PubMed PMID: 38468326; PMCID: PMC10929179.

29. Foy KC, Fisher JL, Lustberg MB, Gray DM, DeGraffinreid CR, Paskett ED. Disparities in breast cancer tumor characteristics, treatment, time to treatment, and survival probability among African American and white women. NPJ Breast Cancer. 2018;4:7. Epub 20180320. doi: 10.1038/s41523-018-0059-5. PubMed PMID: 29582015; PMCID: PMC5861087.

30. Siddharth S, Parida S, Muniraj N, Hercules S, Lim D, Nagalingam A, Wang C, Gyorffy B, Daniel JM, Sharma D. Concomitant activation of GLI1 and Notch1 contributes to racial disparity of human triple negative breast cancer progression. Elife. 2021;10. Epub 20211210. doi: 10.7554/eLife.70729. PubMed PMID: 34889737; PMCID: PMC8664295.

31. Siddharth S, Sharma D. Racial Disparity and Triple-Negative Breast Cancer in African-American Women: A Multifaceted Affair between Obesity, Biology, and Socioeconomic Determinants. Cancers (Basel). 2018;10(12). Epub 20181214. doi: 10.3390/cancers10120514. PubMed PMID: 30558195; PMCID: PMC6316530.

32. Curtis C, Shah SP, Chin SF, Turashvili G, Rueda OM, Dunning MJ, Speed D, Lynch AG, Samarajiwa S, Yuan Y, Gräf S, Ha G, Haffari G, Bashashati A, Russell R, McKinney S, Aparicio S, Brenton JD, Ellis I, Huntsman D, Pinder S, Murphy L, Bardwell H, Ding Z, Jones L, Liu B, Papatheodorou I, Sammut SJ, Wishart G, Chia S, Gelmon K, Speers C, Watson P, Blamey R, Green A, MacMillan D, Rakha E, Gillett C, Grigoriadis A, De Rinaldis E, Tutt A, Parisien M, Troup S, Chan D, Fielding C, Maia AT, McGuire S, Osborne M, Sayalero SM, Spiteri I, Hadfield J, Bell L, Chow K, Gale N, Kovalik M, Ng Y, Prentice L, Tavaré S, Markowetz F, Langerød A, Provenzano E, Purushotham A, Børresen-Dale AL, Caldas C. The genomic and transcriptomic architecture of 2,000 breast tumours reveals novel subgroups. Nature. 2012;486:346–52. doi: 10.1038/nature10983. PubMed PMID: 22522925.

33. Pereira B, Chin SF, Rueda OM, Vollan HK, Provenzano E, Bardwell HA, Pugh M, Jones L, Russell R, Sammut SJ, Tsui DW, Liu B, Dawson SJ, Abraham J, Northen H, Peden JF, Mukherjee A, Turashvili G, Green AR, McKinney S, Oloumi A, Shah S, Rosenfeld N, Murphy L, Bentley DR, Ellis IO, Purushotham A, Pinder SE, Borresen-Dale AL, Earl HM, Pharoah PD, Ross MT, Aparicio S, Caldas C. The somatic mutation profiles of 2,433 breast cancers refines their genomic and transcriptomic landscapes. Nat Commun. 2016;7:11479. Epub 20160510. doi: 10.1038/ncomms11479. PubMed PMID: 27161491; PMCID: PMC4866047.

34. Baghirova S, Hughes BG, Hendzel MJ, Schulz R. Sequential fractionation and isolation of subcellular proteins from tissue or cultured cells. MethodsX. 2015;2:440–5. Epub 20151107. doi: 10.1016/j.mex.2015.11.001. PubMed PMID: 26740924; PMCID: PMC4678309.

